# Local Delivery of Stromal Cell-Derived Factor-1α Improves the Pregnancy Rate of Injured Uterus through the Promotion of Endometrial and Vascular Regeneration

**DOI:** 10.1101/251579

**Authors:** Hongxai Cai, Bingbing Wu, Yixiao Liu, Yu Li, Libing Shi, Lin Gong, Yaxian Xia, BoonChin Heng, Huiling Wu, Hongwei Ouyang, Zhenghua Zhu, Xiaohui Zou

## Abstract

Severe infection and mechanical injury of the uterus may lead to infertility and miscarriage. Currently, there is a lack of effective treatment modality for functional repair of uterine injury. To address this clinical challenge, this study aimed to develop a chemotactic composite scaffold by incorporating recombinant human stromal cell-derived factor-1α (rhSDF-1α) into a silk fibroin-bacterial cellulose (SF-BC) membrane carrier. A rat model of uterine injury was utilized for this study, which was composed of three groups: blank control, implantation with SF-BC only or SF-BC loaded with rhSDF-1α. The tissue regeneration efficacy of the three groups were analyzed and compared. The results showed that SF-BC loaded with rhSDF-1α significantly enhanced endometrial regeneration and arteriogenesis of the injured rat uterus, which led to improved pregnancy outcomes, thus indicating much promise for functional uterine repair and regeneration.

## 1. Introduction

The uterus is an important hormone-responsive organ in female mammals, and is essential for the implantation and development of embryos. However, severe infection and certain serious uterine injuries, such as currettage may cause uterine adhesion, while caesarean section may result in scar formation, infertility, recurrent miscarriages and other obstetrical complications, such as placenta previa and placenta accreta. Therefore, it is essential to repair the damaged uterus to enable a normal pregnancy (Micili *et al.*, 2013). Functional repair and regeneration of the injured/damaged uterus still remains a formidable challenge in clinical practice. Tissue engineering and regenerative medicine could be a promising way to solve this problem.

Migration of endogenous cells to the injury site and adequate vascularization are crucial to tissue regeneration. Stromal cell-derived factor −1α (SDF-1α), also known as CXCL12, is a well-known homing factor for certain stem/progenitor cells. It can recruit tissue resident stem cells such as mesenchymal stem cells (MSCs) via the CXCL12-CXCR4 signaling axis (Son *et al.*, 2006). SDF-1α has already been applied in the regeneration of the liver, heart, skin and nervous system (Imitola *et al.*, 2004; Nakamura *et al.*, 2013; Song *et al.*, 2014). Hence, the delivery of exogenous SDF-1α to the injury site of the uterus might promote tissue regeneration by inducing migration of endogenous cells.

Silk fibroin (SF)–based scaffolds have been widely utilized in bone repair, bladder regeneration and wound healing due to its good biocompatibility, mechanical properties, high plasticity and biodegradability (Tu *et al.*, 2013). Bacterial cellulose (BC) is produced by certain types of bacteria and is characterized by good biocompatibility, high tensile strength, porosity, strong water absorption, making it suitable for tissue engineering of bone and nerve, repair of tympanic membrane perforation and vascular grafts(Gu *et al.*, 2014; Kim *et al.*, 2013). Recently, we have fabricated a composite scaffold of SF and BC that combined the mechanical strength of SF and water absorption properties of BC (Zhu *et al.*, 2015).

In this study, we used the fabricated composite SF-BC membrane as a carrier to incorporate the rhSDF-1α and investigated the effects of the rhSDF-1α functional regeneration of the uterine injury.

## 2. Materials and Methods

### 2.1 Physical and chemical properties of SF-BC membrane

#### 2.1.1. Production of silk fibroin-bacterial cellulose composite scaffold (SF-BC)

The silk fibroin-bacterial cellulose composite scaffold (SF-BC) was prepared according to previous methods (Zhu *et al.*, 2015). Briefly, the silk fibroin was prepared by dissolving degummed silk fiber in CaCl_2_-CH_3_CH_2_OH-H_2_O (1:2:8 in mole ratio) at 60°C for 40 min, and dialyzing against distilled water (Ajisawa 1998; Lindsay *et al.*, 2011). Then, 1g/100ml silk fibroin was added in fermentation medium (50g/L glucose, 10g/L peptone, 1g/L citric acid, 1g/L Na_2_HPO_4_•12H_2_O, 3g/L KH_2_PO_4_, 1.46g/L MgSO_4_•7H_2_O) and cultivated with Activated *Acetobacter Xylinum* (1.1812), purchased from the Institute of Microbiology Chinese Academy of Science, at 30°C for 10 days without shaking the incubator. The SF-BC membrane thus obtained were first washed in distilled water to remove fermentation medium, then immersed in 1% (w/v) NaOH solution at 80°C for 3h, with several rinses in distilled water every hour. Subsequently, 1% (w/v) NaOH solution was neutralized by 0.1% acetic acid, and the SF-BC membrane was then rinsed with distilled water, followed by lyophilization, then SF-BC was sterilized by autoclaving (Figure 1a).

**Figure 1:**
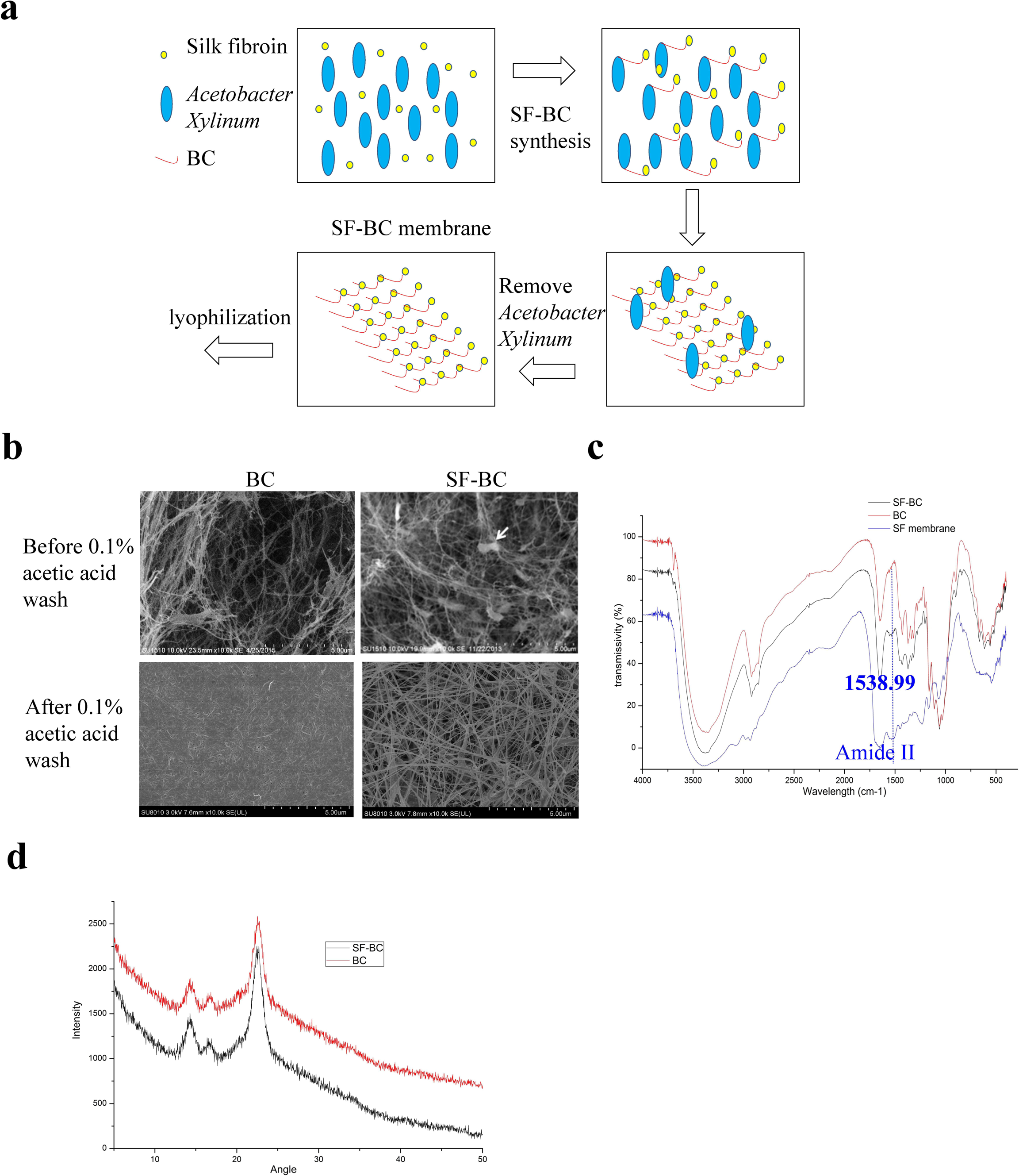
Physical and chemical analysis of the SF-BC membrane. (a) Schematic diagram of SF-BC membrane formation. Acetobacter Xylinum was cultured in fermentation medium containing SF. In the process of BC formation, SF is combined with BC, to form the SF-BC membrane. (b) SEM images of BC and SF-BC showed more porous structure within the SF-BC membrane. (c) FTIR analysis of SF-BC showed a peak at 1538.99 cm^-1^ that is characteristic of the amide II group of the silk component. (d) XRD analysis of the SF-BC membrane showed three peaks characteristics of the BC crystal.

#### 2.1.2. Scanning electron microscopy (SEM) imaging of the SF-BC membrane

The BC and SF-BC membrane without cells were mounted on aluminum stubs and coated with gold directly. Human uterine cells cultured on SF-BC membrane for 7 days were fixed with 0.25% glutaraldehyde solution for 24h. After rinsing with PBS for 3 times, the samples were soaked in OsO_4_ (Ted Pella) for 1h, and washed again three times with PBS. The samples were then dehydrated with a concentration gradient of acetone (30%, 50%, 70%, 80%, 90%, 95%, 100%, v/v), followed by drying. Lastly the samples were placed on aluminum stubs and coated with gold. The cells cultured on the SF-BC membrane were visualized by scanning electron microscopy (Hitachi SU-8010N, Japan) at an accelerating voltage of 15kV.

#### 2.1.3. Fourier transforms infrared spectroscopy (FT-IR) of the SF-BC membrane

The BC, SF and SF-BC membranes were air dried, and were subjected to fourier transform infrared (FT-IR) spectroscopy with a Nicolet AVA TAR370 spectrometer (Thermo Fisher Scientific, USA). The scanning range was from 4000 to 400 cm^-1^ with a resolution of 4 cm^-1^.

#### 2.1.4. X-Ray Diffraction (XRD) analysis of the SF-BC membrane

The SF-BC and BC membranes were air dried, and X-Ray Diffraction (XRD) analysis was measured with an x-ray diffractometer at 2.7KW (Shimadzu XRD-6000, Japan), with the tested angle being from 0° to 40°.

#### 2.1.5. Mechanical properties of the SF-BC membrane

Mechanical properties of BC and SF-BC membrane were tested by using an Instron tension/compression system with Fast-Track software (Model 5543, Instron, Canton, MA, USA) under hydrated condition. The membranes were cut into a rectangular piece of 4 mm in width and 13 mm in length, after saturated with 0.9% saline for 12 hours at 37 °C, 2mm in length at both sides were clamped by hemostatic forceps. The membranes were then stretched at a speed of 5 mm/min. At least 4 replicate samples of each membrane were measured, data were presented as Mean ± SEM (Standard Error of Mean). The experiment was performed at room temperature.

### 2.2 Biological properties of the SF-BC membrane

#### 2.2.1. Cytotoxicity evaluation of the SF-BC membrane

SF-BC was sterilized by autoclaving, a 2cm×1cm SF-BC membrane section was immersed in 0.67 ml of high-glucose DMEM (DMEM/H) containing 10%(v/v) FBS for l of DMEM/H containing 1000 L929 cells (Chinese Academy of Sciences, Beijing, China) were seeded in each well of a 96-well plate. After 24h, the culture medium was replaced by media containing 5% (v/v) of extracts from the membrane, fresh DMEM/H containing 10% (v/v) FBS with 1% (w/v) penicillin and 1% (w/v) penicillin-streptomycin, within a 5% CO_2_ incubator set at 37 °C for 48h. And 100μl of DMEM/H containing 1000 L929 cells (Chinese Academy of Sciences, Beijing, China) were seeded in each well of a 96-well plate. After 24h, the culture medium was replaced by media containing 5% (v/v) of extracts from the membrane, fresh DMEM/H containing 10% (v/v) FBS with 1% (w/v) penicillin streptomycin, 5% (v/v) DMSO, and cultured for 3 days without changing the medium. The cytotoxicity was then assessed by comparing the OD value at 450 nm with the Cell Counting KIT-8 (CCK-8, Dojindo, Kumamoto, Japan). The OD value is positively correlated with cell number.

#### 2.2.2. Cell proliferation assay on the SF-BC membrane

Human endometrial cells were isolated from the normal portions of the uterus of leiomyoma patients from the First Affiliated Hospital, School of Medicine, Zhejiang University. Approval for utilizing the patient samples in this study was obtained from Ethics Committee of the First Affiliated Hospital, School of Medicine, Zhejiang University. Rat uterine cells were prepared from postnatal rat uterus purchased from the Zhejiang Academy of Medical Science. All experimental procedures involving animals were approved by the Ethics Review Board for Animal Studies of Zhejiang University. Human and rat uterine cells were prepared by mincing the tissues for 5-7 min using a pair of small surgical scissors followed by digestion with type I collagenase (Gibco) in DMEM/F12overnight at 4□. The digested tissues were triturated into individual cell suspensions by a 1 ml pipette. Human endometrial cells were seeded on SF-BC membrane at a density of 3000cells/well, and the OD value at 450 nm was measured with CCK8 after 1, 3, 5 and 7 days of culture. Increase in absorbance values at 450 nm corresponded to the proliferation of human uterine cells.

#### 2.2.3. Cell morphology on the SF-BC membrane

Four thousand human endometrial cells were seeded on the SF-BC membrane in a 96-well plate, and were cultured in a 5% CO_2_ incubator at 37 °C with regular changes in culture medium every 2-3 days. After 7 days, the culture medium was removed, and cells were washed three times with PBS, followed by fixation in 4% (w/v) paraformaldehyde for 30 min, and subsequent rinsing with PBS for another 3 times. Next, we observed the cytoskeleton and nuclei by immunofluorescence staining of F-actin which was detected using TRITC-conjugated Phalloidin (1:1500, Millipore, USA) and DAPI (1:1200, Beyotime Institute of Biotechnology, China) separately. The cell morphology was imaged under confocal microscopy (Olympus, BX61W1-FV1000, Japan).

### 2.3 Migration of rat uterine cells *in vitro* under the influence of SDF1α

The effects of rhSDF-1α on the migration of rat uterine cells were assessed by transwell assay (membrane pore size 8 mm, Costar 3422) as described previously (Shen *et al.*, 2010). Rat uterine cells (P_0_) were trypsinized and seeded onto the upper chamber of transwell inserts at a density of 1×10^4^/well within 100μl of serum-free DMEM/F12 per well of a 24-well plate. Then 600μl of serum-free DMEM/F12 or 600μl of serum-free DMEM/F12 containing 100ng/ml rhSDF-1α (R&D Systems, Minneapolis, MN, USA) were added into the lower chamber to induce migration of rat uterine cells (Zhang *et al.*, 2013; Miyazaki and Maruyama, 2014).The cells were cultured in a 5% CO_2_ incubator at a temperature of 37 °C for 24h.

In order to confirm the effects of rhSDF-1α, rat uterine cells were pre-treated with 200 ng/ml of ADM3100 (sigma), which is an antagonist of SDF-1α, for 30 min in a 5% CO_2_ incubator at 37°C. Then, 100μl of serum-free DMEM/F12 containing 1×10^4^ pre-treated cells were seeded on the upper chamber of the transwell (Corning), and 600μl of serum-free DMEM/F12 with 100 ng/ml rhSDF-1α in a 5% CO_2_ incubator at 37°C. Then, 100 was added to the lower chamber of the transwell. In another group, the cells seeded on the upper chamber of transwells were cultured in 100μl of serum-free DMEM/F12 with 100ng/ml rhSDF-1α, and 600μl of serum-free DMEM/ F12 with 100ng/ml rhSDF-1α was added to the lower chamber as well.

After 24h, the transwell inserts were taken out from 24-well plates and rinsed three times with PBS, and were fixed with 4% (w/v) paraformaldehyde for 20 min, washed another three times with PBS, and subjected to nuclear staining with DAPI (1:2000, Beyotime Institute of Biotechnology Inc, China) for 10 min. The cells on the upper chamber were scraped to remove adherent cells. The number of cells that migrated to the lower chamber of the transwell was counted within 5 fields (× 100) under microscopy.

### 2.4. Rat uterine horn injury/regeneration model

Twenty-nine female Sprague Dawley (SD) rats (purchased from the Zhejiang Academy of Medical Science) weighing between 200g and 250g were selected, and the 58 uterine horns of the rats were divided into 3 groups, injury alone group (n = 16), SF-BC group (n = 21), and (SF-BC / rhSDF-1α) group (n = 21). All the rats were kept in a specific pathogen-free air-conditioned room and were allowed free access to food and water at the Animal Center of Zhejiang University of Medicine. All experiments were approved by the Ethics Review Board for Animal Studies of Zhejiang University. In order to deliver rhSDF1α, the SF-BC membrane was soaked in a solution of rhSDF1α- at a concentration of 100ng/ml within an incubator set at 37 °C with 5% CO_2_ for 12h according to previous method (Zhang *et al.*, 2013).

The procedure was partly based on the full-thickness injury model of rat uterus that was described previously (Taveau *et al.*, 2004; Yan *et al.*, 2013; Gu *et al.*, 2014; Miyazaki and Maruyama, 2014). After the rats were anesthetized by intra-peritoneal injection of chloral dehydrate (0.4 g/kg), a midline incision in the abdomen was made, and the uterus was exposed. In the injury alone group, a full-thickness defect was made by excising a section that was about 1.0 cm in length and 0.5 cm in width in each uterine horn (Figure 3a), with the mesenterium being retained. The four margins of the injury were marked by 6-0 non-absorbable silk suture. In the other two groups, the SF-BC membrane or SF-BC / rhSDF-1α, the SF-BC membrane was soaked in a solution of rhSDF-1α membrane that was equal to the size of the uterine defect were sutured at the injury site using 7-0 absorbable proline suture via interrupted sutures. SF-BC membranes were either immersed in high-glucose DMEM (DMEM/H) alone or DMEM/H with SDF (100ng/ml) at 37 °C overnight before operation. After operation of the uterus, the abdominal cavity was washed with 0.9% (w/v) normal saline. Then the rectus abdominis was sutured by 6-0 non-absorbable silk suture with continuous suture, and the skin and fascia were sutured by 4-0 non-absorbable silk suture through interrupted suture. During the operation procedure, a piece of gauze, which was soaked in 0.9% (w/v) normal saline, was used to cover the uterus and viscera to avoid drying. All the operated rats were overnight treated with intramuscular injection of penicillin (80000U per rat) continuously for 3 days.

### 2.5. Histological analysis

#### 2.5.1. Hematoxylin and eosin staining

The rats were sacrificed at 30 and 90days after surgery. The surgery site of each uterine horn (injury alone group: n = 4, SF-BC group: n = 6, SF-BC / rhSDF-1α group: n = 6) were fixed in 4% paraformaldehyde, dehydrated with an ethanol gradient, followed by paraffin embedment and sectioning at 5 μm thickness. Then the paraffin sections were stained with hematoxylin and eosin (H&E), and the thickness of the endometrium which starts from the luminal epithelium to the muscle layer at the injury site was measured by Image-Pro Plus software (version 6.0)(Figure 3h).

#### 2.5.2. Immuno-histochemical staining

Sections were dewaxed, rehydrated, and subjected to antigen retrieval treatment with citric acid buffer (PH = 6.0) or pepsin, followed by washing three times with PBS. These were then incubated in 3% (v/v) H_2_O_2_ in methanol for 10min to inactivate endogenous peroxidase. After rinsing with PBS, the samples were treated with blocking solution (1% w/v BSA) for 30 min, and then incubated with primary antibodies at 4 °C overnight. For indirect staining, the slides were further incubated in horseradish peroxidase-conjugated goat anti-mouse immunoglobulin G (IgG) in 1% (w/v) BSA for 2h at room temperature, and then washed 3 times with PBS. The samples were then visualized by the DAB method, and counterstained with hematoxylin solution. The primary antibodies utilized were as follows: monoclonal anti-vimentin (1:50, Dakocytomation, USA), mouse monoclonal anti-pan-cytokeratin (1:200, Abcam, USA), monoclonal anti-smooth muscle actin (1:100, SMA, Boster, China), and the appropriate secondary antibodies (1:50, Invitrogen, China). All the procedures were carried out according to the manufacturer’s instructions.

#### 2.6. Functional test of the regenerated endometrium

In order to determine whether the regenerated endometrium at the surgery site is receptive to the implantation of embryos and can support subsequent embryo development (Taveau *et al.*, 2004; Yan *et al.*, 2013; Gu *et al.*, 2014; Miyazaki and Maruyama, 2014), 13 rats (number of uterine horns in each group: injury alone group, n = 8; SF-BC group, n = 9; SF-BC / rhSDF-1α group, n = 9) were mated with males, and the conception was confirmed if a large number of sperm was visible in the vagina the next day after mating. Then the rats were euthanized at late gestation (19-21 days), and pregnancy rate at the surgery site together with the number of fetuses in each uterine horn were recorded and compared.

#### 2.7. Statistical analysis

The thickness of fibrous tissues around the implanted materials and the distance of cells migrated into the materials were measured, and compared using paired T test in Prism 5.0 software. The number of cells that have migrated to the lower chamber of the transwell inserts, endometrium thickness, number of arteries and number of fetuses in each uterine horn were compared using one-way ANOVA with Bonferroni’s Multiple Comparison Test in Prism 5.0 software. The percentage of endometrial stromal cells, pregnancy rate, and pregnancy rate at the surgery site were analyzed by the X^2^ test and Fisher exact test for categorized variables in SPSS software (Version 19.0, SPSS Inc, Chicago, IL, USA), and all P values less than 0.05 were considered statistically significant.

### 3. Results

#### 3.1. Physical and chemical properties of the SF-BC membrane

The fabricated SF-BC membrane was imaged under digital camera and SEM. Digital camera images showed that the SF-BC membrane was thin and white in color (Figure 2e). SEM imaging revealed that SF-BC had a more porous structure compared to that of BC, with the fiber diameter at nano-scale (Figure1b). FTIR results showed a specific band from SF-BC at 1538.99 cm^-1^ that was absent in the BC membrane, which might be characteristic of the amide II group of the silk fibroin (Figure1c). XRD analysis of both the BC and SF-BC membrane showed three peaks (BC: 14.29°, 16.83°, 22.51° ; SF-BC:14.28⪚, 16.38°, 22.45°) that were characteristic of the BC crystal (Figure1d). Table 1 presents the mechanical properties of the BC and SF-BC membranes, and the results indicated that the addition of SF (1%) could significantly increase the mechanical properties (maximum load, stiffness and Young’s modulus) of the SF-BC membranes compared with that of the BC membrane. These results showed that the SF-BC membrane has a porous structure with favorable mechanical properties.

**Figure 2:**
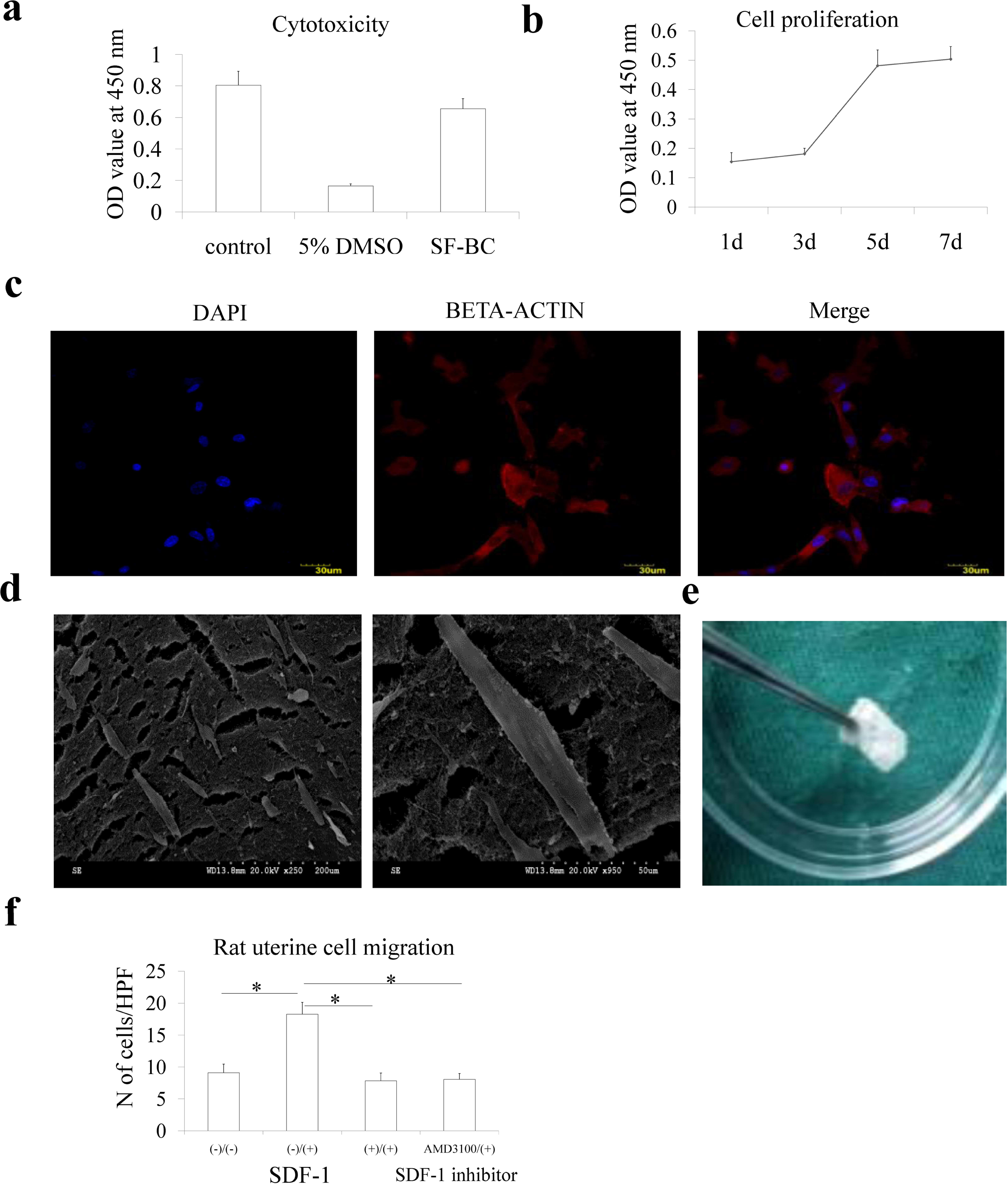
Biological properties of the SF-BC membrane. (a) Cytotoxicity evaluation of the SF-BC membrane. (b) Cell proliferation assay. (c) Confocal imaging of human uterine cells cultured on the SF-BC membrane for 7d. (d) SEM imaging of human uterine cell morphology cultured on the SF-BC membrane (e) External structure of the SF-BC membrane. (f) Chemotactic effects of rhSDF-1α on the migration of rat uterine cells. (-)(-):culture medium in the upper and lower chambers of the transwell inserts were serum-free DMEM/F12; (-)(+): culture medium in the upper chamber was serum-free DMEM/F12, and lower chamber was serum-free DMEM/F12 with rhSDF-1α (100ng/ml); AMD3100/(+):uterine cells were pretreated with AMD3100 for 20 min, and the culture medium was serum-free DMEM/F12 in the upper chamber, while that in the lower chamber was serum-free DMEM/F12 with rhSDF-1α (100ng/ml). *:p. value < 0.05. HPF: high power field. (100ng/ml); (+)(+): both upper and lower chambers were serum-free DMEM/F12 with rhSDF-1α 1α(100ng/ml). *:p. value < 0.05. HPF: high power field.

**Table 1:** Mechanical properties of BC and SF-BC composite membranes.

| Membrane | Max load (N) | Stiffness (N/m) | Young's modulus (Mpa) |
| --- | --- | --- | --- |
| BC(n=5) | 1.827±0.382<sup>a</sup> | 0.963 ±0.144<sup>a</sup> | 2.911±0.437<sup>a</sup> |
| 1%SF-BC(n=6) | 2.435±0.246<sup>b</sup> | 2.782±0.617<sup>b</sup> | 8.411±1.864<sup>b</sup> |

Data were presented as Mean ± SEM (Standard Error of Mean), difference in the letter (a & b) indicating the significant difference between two groups(p. value < 0.05).

#### 3.2. Biological properties of the SF-BC membrane

Additionally, the cytotoxicity test (Figure2a) and cell proliferation assay (Figure2b) showed that the SF-BC membrane has good biocompatiblity. In order to determine whether the SF-BC membrane affected cell morphology, human endometrial cells were seeded on it for 7 days. The confocal microscopy images showed that the cells adopted various shapes, with the cytoskeleton being stretched out on the membrane (Figure2c).This was also supported by the SEM images showing the cells being spread out on the membrane surface (Figure2d). These results thus suggest that the SF-BC membrane was biocompatible and conducive for uterine cells, which is of utmost importance for tissue regeneration at the injury site.

#### 3.3. The effects of rhSDF-1α on rat uterine cells

The chemotactic effects of rhSDF-1α on rat uterine cells were tested by transwell assay *in vitro*, and the results showed that the number of cells that migrated through the upper chamber (in which rhSDF-1α was added) into the lower chamber was significantly more than that of the blank (18.27 ± 1.87 vs 9.09 ± 1.38, p < 0.05). This effect was blocked by adding rhSDF-1α to the lower chamber, or pre-treating all of the cells with AMD3100 (an antagonist of SDF-1) (7.85 ±1.21 vs 9.09 ± 1.38, 8.06 ± 0.94 vs 9.09 ± 1.38, respectively, both p > 0.05) (Figure2f). These results thus demonstrate that rhSDF-1α could promote the migration of rat uterine cells *in vitro*.

#### 3.4. Regeneration of rat uterus treated with rhSDF-1α after surgery loaded SF-BC membrane after surgery

Thirty days after surgery, all of the uterine horns kept patency. The uterine horns implanted with SF-BC membrane showed only slight adhesion at the injury site. However, certain uterine horns in the injury alone group exhibited severe adhesion to the viscera, particularly to the small intestine. Moreover, there were fewer blood vessels in the injury site alone group. Although the SF-BC membrane degraded slowly, it interacted well with the adjacent tissues (Figure 3b).Uterine horns implanted with the SF-BC membrane retained their original shape at 90 days post-operation, while uterine horns in the injury alone group exhibited stenosis at the injury site (Figure 3c), which may have an adverse effect on the pregnancy outcomes.

**Figure 3:**
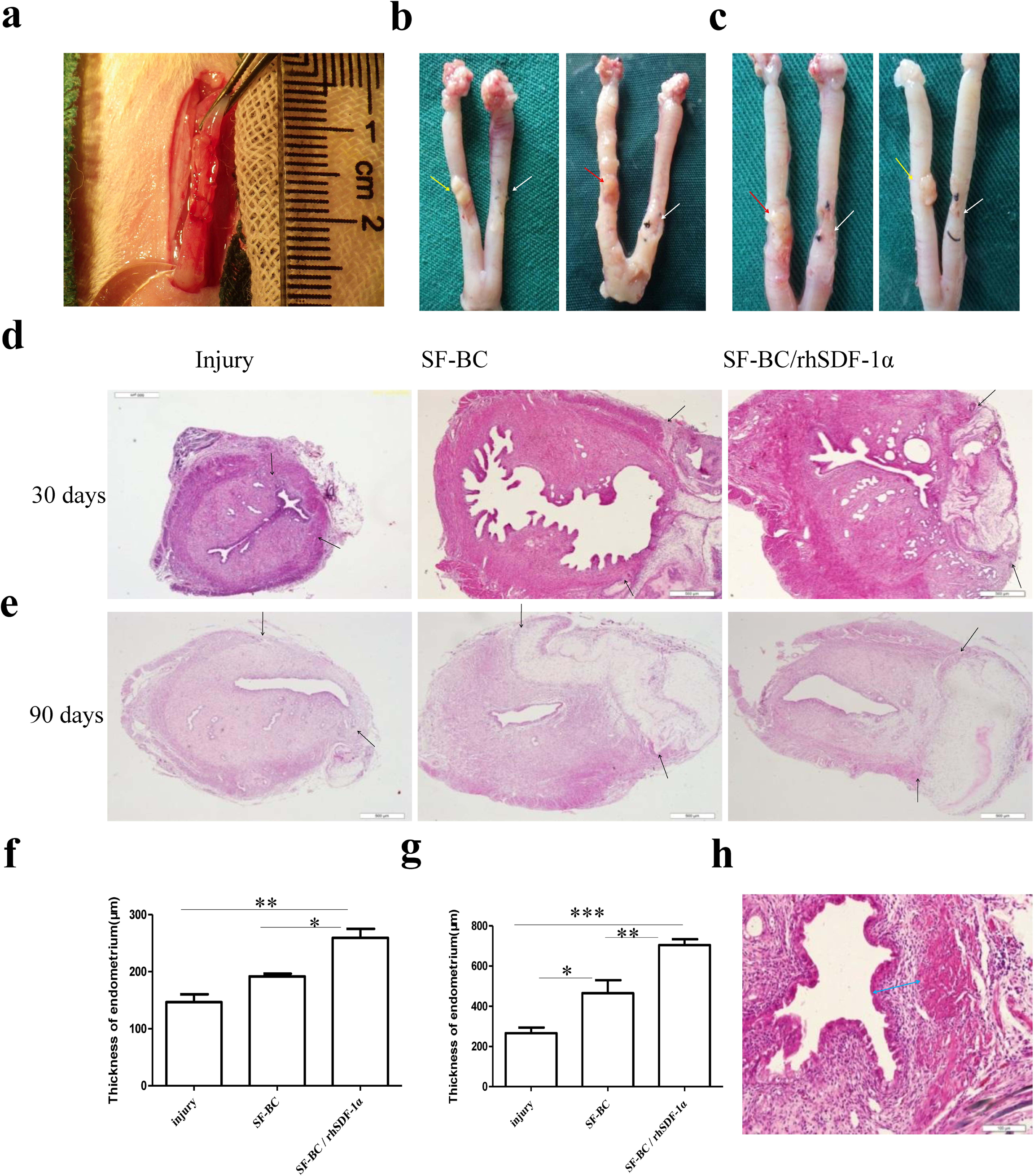
Regeneration of rat uterus treated with the SF-BC membrane loaded with rhSDF-1α. (a) Rat uterus implanted with the SF-BC membrane. Gross appearance of rat uterus at 1 month (b) and 3 months (c) after surgery, white arrows: injury alone group; yellow arrows: SF-BC group; red arrows: SF-BC with rhSDF-1α and (e) represent H&E staining of the regenerated endometrium at 30 days and 90 group. (d) days post-surgery, separately. Scale bars = 500 μm. (f) and (g) compare the thickness of endometrium formed at the injury site at 30 days and 90 days post-surgery, respectively. (h) Measurements of endometrial thickness. Arrows indicate the range of injury. *: p. value < 0.05; **: p. value < 0.01; ***: p. value < 0.001.

Histological analysis showed that the regenerated endometrium at the surgery site had intact luminal epithelium at 30 days post-operation, with secretory glands appearing at the injury site in both the SF-BC and SF-BC/rhSDF-1α groups, whereas only a few glands could be observed in the injury alone group (Figure 3d). Furthermore, the endometrium in the SF-BC/rhSDF-1α group was thicker than that of both the SF-BC group and the injury alone group (259.29 ± 15.94μm vs 191.55 ± 4.79μm, 259.29 ± 15.94μm; vs 146.75 ± 13.92μm, respectively), but there was no statistically significant difference in thickness of the endometrium between the SF-BC group and the injury alone group (Figure 3f).

Ninety days after surgery, more secretory glands had regenerated at the injury site in both the SF-BC and the SF-BC/rhSDF-1α groups, whereas there were only a few glands present in the injury alone group(Figure 3e). Although the endometrium thickened in all groups, the endometrium in the SF-BC/rhSDF-1α group was thicker than that of both the SF-BC group and the injury alone group (704.00 ± 29.52μm 465.40 ± 65.14μm, 704.00 ± vs 266.08 ± 28.56μm, respectively). The endometrium in the SF-BC group was thicker than that of the injury alone group (465.40 ± 65.14μm vs 266.08 ± 28.56μm) (Figure 3g). In conclusion, the results showed that the SF-BC membrane could promote regeneration of the endometrium.

#### 3.5 Pregnancy outcomes of the regenerated endometrium

One of the most important functions of the uterus is to receive implantation of embryos and support subsequent development of the fetus. In this study, we found that some regenerated endometrium received implantation of embryos in all 3 groups at 30 days post-injury (Figure 4). In the injury alone group, 4/8 of them had fetuses in the uterine horns, which was lower than that in the SF-BC group (7/9) and the SF-BC/rhSDF-1α group (8/9). However, there was no statistically significant difference between them. Moreover, 2/8 uterine horns had an embryo implanted at the surgery site of the injury alone group, which was less than that in the SF-BC group (5/9) and the SF-BC/rhSDF-1α group (5/9), but the observed differences were not statistically significant, probably due to the small sample size(Table 2).

**Figure 4:**
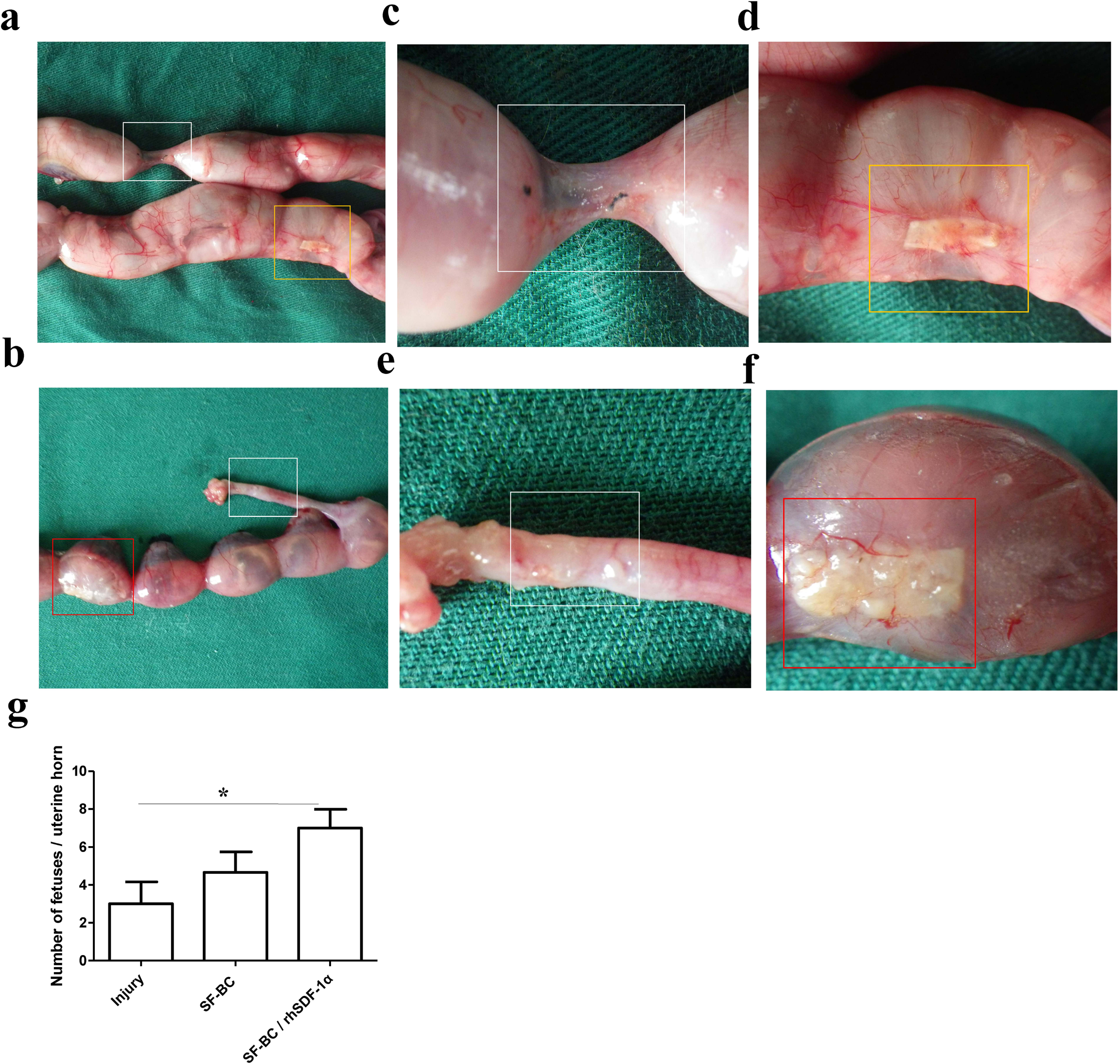
Pregnancy outcomes of injured uterus at 30 days post-surgery. (a) and (b) showdeveloping fetuses at the injury site. (c) and (d) representenlarged magnification of the frames in (a). (e) and (f) represent magnification of the frames in (b).(g) Number of fetuses per uterine horn. White frames indicate injury alone site, yellow frames indicate injury site transplanted with only SF-BC, and red frames indicate injury site implanted with SF-BC loaded with rhSDF-1α

**Table 2:** Pregnancy rates, numbers of fetuses/uterine horn and pregnancyrates at the injury site after 19-20 days post-mating with male rats.

|  | **Injury(n=8)** | **SF-BC(n=9)** | **SF-BC/rhSDF-1α(n=9)** |
| --- | --- | --- | --- |
| **Pregnancy rate** | 4/8 | 7/9 | 8/9 |
| **Number of fetuses per uterine horn** | 3.0 ± 1.2 (0-7)<sup>a</sup> | 4.4 ± 1.0 (0-7)<sup>ab</sup> | 6.8 ± 1.0 (0-10)<sup>b</sup> |
| **Pregnancy rate at surgery site** | 2/8 | 5/9 | 5/9 |

Number of fetuses per uterine horn: Data were presented as Mean ± SEM (Standard Error of Mean), difference in the letter (a & b) indicating the significant difference between two groups (p. value < 0.05).

In addition, there was variation in the number of fetuses developed in each uterine horn. In the SF-BC/rhSDF-1α group, the number of fetuses developed per uterine horn (7.00 ± 0.99) was more than that in the injury alone group (3.00 ± 1.16). But there was no statistically significant difference between the SF-BC group (4.67 ±1.08) and the SF-BC/rhSDF-1α group, as well as between the injury alone group and the SF-BC group (Table 2). This data (Figure 4g) suggests that SF-BC/rhSDF-1α improved pregnancy outcomes at 30 days after surgery.

### 3.6. Mechanisms of the effects of SF-BC/rhSDF-1α on the functional regeneration of the uterus

#### 3.6.1. Recruitment of endometrial stromal cells at the injury site

In order to determine the proportion of endometrial stromal cells within the regenerated endometrium, sections of the endometria were stained with an anti-vimentin antibody. The percentage of endometrial stromal cells in the SF-BC/rhSDF-1α group was higher than that in the injury alone group at 30 days post-surgery (45.11 ± 2.27% vs22.25 ± 4.25%) (Figure 5a, Table 3). However, there were no statistically significant differences between the SF-BC group and the injury alone group, as well as between the SF-BC/rhSDF-1α group and the SF-BC group (34.95 ± 3.98% vs 22.25 ± 4.25%, 45.11 ± 2.27% vs 34.95 ± 3.98%). Figure 5b showed that there were no statistically significant differences in the percentages of endometrial stromal cells amongst the three groups (SF-BC/rhSDF-1α group: 34.95 ± 3.98%, SF-BC group: 37.64 ± 3.02%, injury alone group: 37.04 ± 2.82%) at 90 days post-injury (Table 3). This may be due to the proliferation of stromal cells at the injury site and migration of stromal cells from the adjacent normal site. These data suggests that SF-BC/rhSDF-1α play a role in the migration of endometrial stromal cells at the surgery site within a short duration.

**Figure5:**
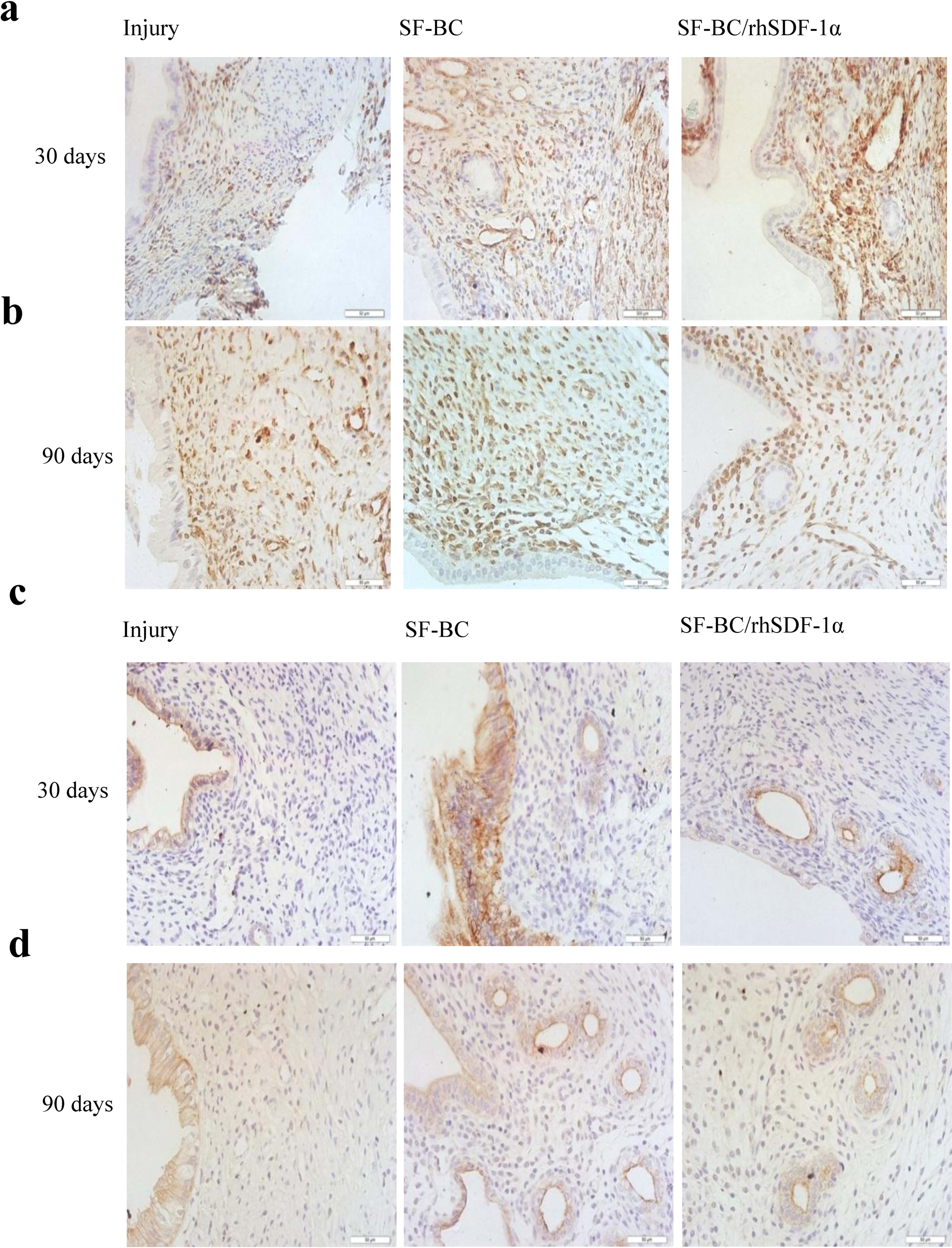
Characterization of regenerated endometria of the rat uterus. Immunohistochemical staining of endometrial stromal cells (a) and (b) shows endometrial stromal cells in the newly formed endometrium: immunohistochemical staining of vimentin at 30 days and 90 days post-surgery, respectively. Scale bars = 50 μm. (c) and (d) show the presence of endometrial luminal and glandular epithelial cells by immunohistochemical staining of pan-cytokeratin at 30 days and 90 days post-surgery, respectively. Scale bars = 50 μm.

**Table 3:**
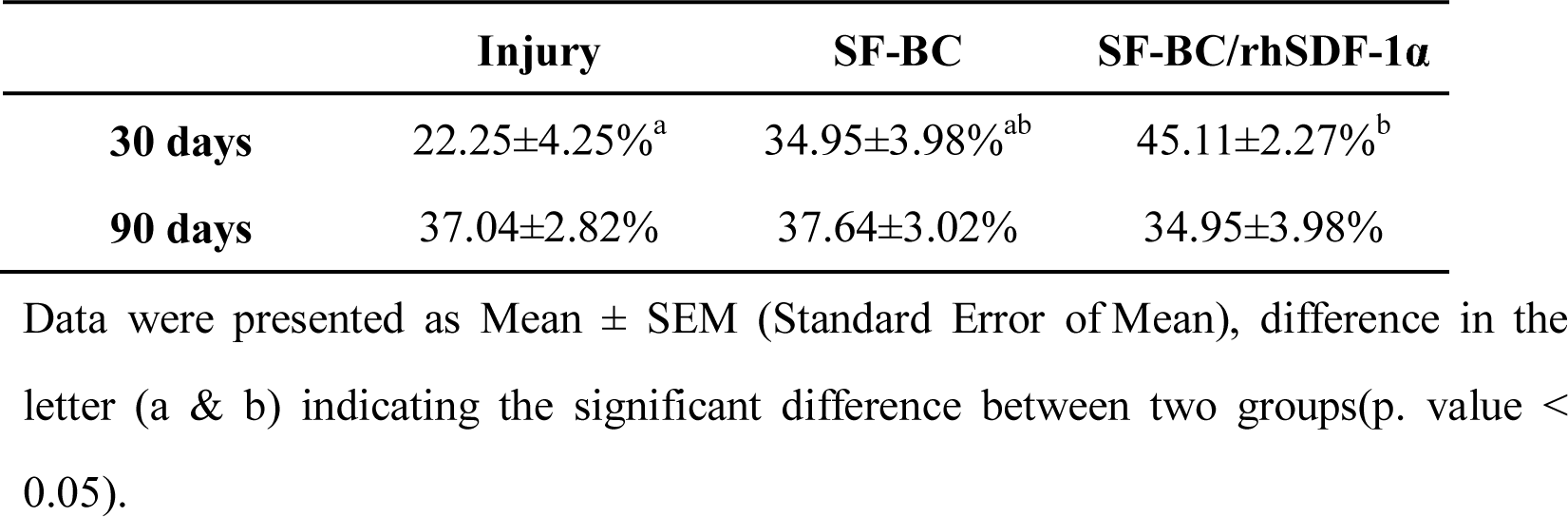
Percentage of vimentin-positive cells amongst the three groups.

#### 3.6.2. Formation of endometrial epithelial cells at the injury site

Endometrial epithelial cells can be classified into two distinct types: luminal epithelial cells and glandular epithelial cells. The latter plays an important role in decidualization, implantation and embryo development. After the sections were labeled with an anti-pan-cytokeratin antibody, luminal epithelial cells could be detected in the injury site among the three groups at 30 days post-surgery. However, significant numbers of glandular epithelial cells were present only in the SF-BC group and the SF-BC/rhSDF-1α group, while few of these were present in the injury alone group (Figure 5c).

Ninety days after injury, more glandular epithelial cells were detected at the surgery site of the SF-BC group and the SF-BC/rhSDF-1α group. Nevertheless, the number of glandular epithelial cells in the injury alone group at 90 days post-surgery was similar to that at30 days post-surgery (Figure 5d). This data suggests that the SF-BC membrane facilitated the formation of endometrial secretory glands, and this may contribute to differences in pregnancy outcomes amongst the 3 groups. The lack of secretory glands in the regenerated endometrium of the injury alone group would result in endometrial dysfunction, which may affect subsequent pregnancy outcomes.

#### 3.6.3. Arteriogenesis in the regenerative endometrium

Arteriogenesis is crucial in tissue repair as the rich blood supply not only provides nutrients, but also various types of growth factors, which can modulate cell migration, proliferation and differentiation. To investigate whether rhSDF-1α could enhance arteriogenesis, the sections were stained with an anti-α-SMA antibody. The number of arteries was estimated by counting α-SMA positive blood vessels in 5 HPF.

At 30 days post-surgery, the number of arteries in the SF-BC/rhSDF-1α group was more than that in the injury alone group and the SF-BC group (7.45 ± 0.38 vs 1.13 ± 0.31, 7.45 ± 0.38 vs 2.20 ± 0.20, separately) (Figure 6a, 6c). There was no significant difference between the SF-BC group and the injury alone group (2.20 ± 0.20 vs 1.13 ± 0.31) (Figure 6a, 6c). At 90 days post-surgery, the number of arteries in the injury alone group was less than that in the SF-BC group and SF-BC/rhSDF-1α groups (3.03 ± 0.41 vs 6.65 ± 0.40, 3.03 ± 0.41 vs 5.50 ± 0.23, respectively). (Figure 6b, 6d) No significant difference in the number of arteries was found between the SF-BC group and the SF-BC/rhSDF-1α demonstrated that SF-BC/rhSDF-1α -SMA positive blood vessels in 5 -SMA antibody. The group (Figure 6b, 6d). The data thus demonstrated that SF-BC/rhSDF-1μ promoted arteriogenesis at the injury site within a short time duration, and that the SF-BC membrane was beneficial for the formation of arteries at the injury site over a longer duration.

**Figure 6:**
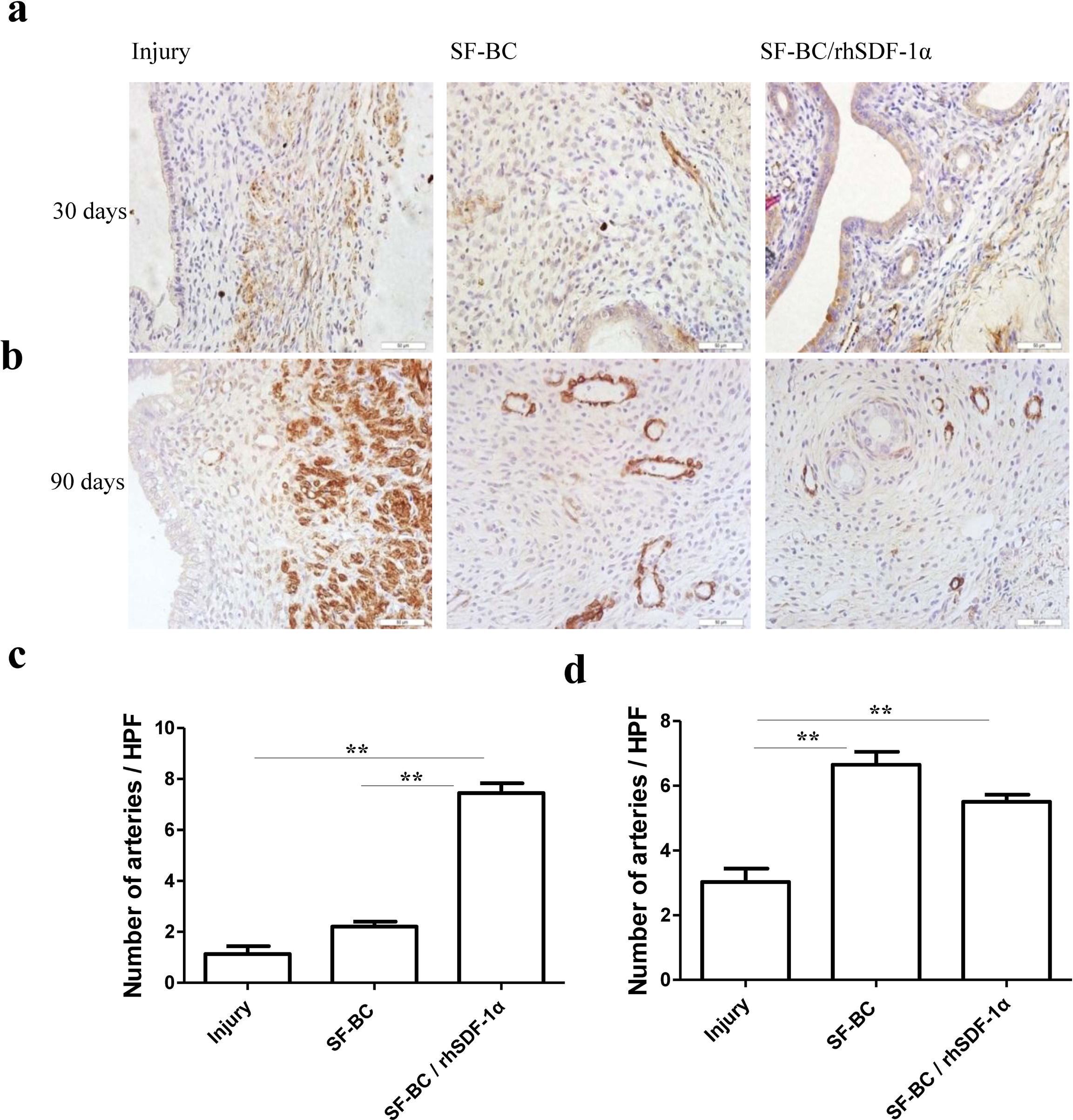
Immunohistochemical staining of arteries at the injury site. (a) and (b) reveal formation of endometrial arteries in the injury sites at 30 days and 90 days post-surgery by immunohistochemical staining of α-SMA, separately. (c) Numbers of arteries formed in the injury site amongst the 3 groups at 30 days post injury, and (d) Number of arteries at 90 days post-injury. Scale bars = 50 μm. **: p. value < 0.01.

## 4. Discussion

Curettage, infections and cesarean section may lead to amenorrhea, infertility, and pregnancy complications (Micili *et al.*, 2013). In recent years, tissue engineering has been applied to address this clinical challenge. Some studies attempted to utilize SIS to reconstruct the injured uterus, however, SIS collapsed or became twisted when it is longer than 1cm (Taveau *et al.*, 2004). Miyazaki and Maruyama (2014) found that decellularized uterine matrix demonstrated some efficacy in promoting uterine regeneration (Miyazaki and Maruyama, 2014). Nevertheless, human uterine tissue is not readily available due to ethical and legal issues, which limits the application of the aforementioned scaffold. Because the uterine cavity in humans needs to expand about 1000 times during late gestation, scaffolds suitable for uterine repair should have good biocompatibility and excellent mechanical properties, besides being readily and easily accessible. In this study, we used a composite scaffold of silk fibroin and bacterial cellulose which can fulfill these requirements, and could act as a good carrier for the delivery of biological factors.

Silk fibroin and bacterial cellulose are two biological materials widely utilized in scaffold fabrication and tissue engineering (Kim *et al.*, 2013; Tu *et al.*, 2013). We fabricated the composite scaffold of SF and BC to synergize the strength of SF with the elasticity and water absorption properties of BC (Zhu *et al.*, 2015). In the process of fabricating the composite membrane scaffold (Figure 1), the bacterial cellulose and silk fibroin were fermented in a strongly acidic environment (pH<3). This resulted in intermolecular dehydration, in situ polymerization between cellulosic residues (-OH) and silk fibroin residues (-COOH and -NH2), and the binding of silk fibroin molecules to the bacterial cellulose, which resulted in pore formation within the composite membrane scaffold (Figure1b). The porous interior of the composite membrane scaffold made it easier for somatic cells to get access to the internal space and grow. With the gradual degradation of SF upon implantation *in vivo*, the space between the fibers increased, which could further facilitate the growth and differentiation of cells. In the SF-BC composite membrane, the amino acid constituents of SF could facilitate the aggregation of collagen that would make it conducive for cell growth and differentiation in the regenerating uterus (Xu *et al.*, 2012). These results could explain the proliferation of uterine cells *in vitro* (Figure 2b) and the increase in proportion of vimentin-positive stromal cells detected in the SF-BC composite membrane group *in vivo* (Figure 5). The SF-BC composite membrane fabricated in this study was safe and biocompatible with negligible cytotoxicity and could support the proliferation of uterine cells (Figure 2b). Because the plasticity and tensile strength of the composite membrane scaffold was similar to that of the rat uterus, this could promote regeneration of the uterus after injury and physically expand to support the growth of implanted embryos. The structure of the composite SF-BC membrane was also favorable to the loading of exogenous factors, which could further promote repair and regeneration of injured tissues.

Stromal cell-derived factor-1α (SDF-1α), also known as CXCL12, is a homing factor for certain stem/progenitor cells. In this study, we found that rhSDF-1α (SDF-1α could promote the migration of uterine cells *in vitro*, as well as enhance regeneration of the injured endometrium *in vivo*, leading to improved pregnancy outcome after uterine injury at 30 days post-surgery. In contrast, uterine repair with other factors e.g. bFGF, VEGF could only result in blastocyst implantation at about 90 days post-surgery, as reported by previous studies (Lin *et al.*, 2012; Ding *et al.*, 2014). The data thus suggests that rhSDF-1α was more efficacious and could hasten regeneration and pregnancy outcomes 60 days earlier, as compared to previous studies. Additionally, the number of fetuses per uterine horn (1.3 ± 0.5) in post-injury uterus that was partially reconstructed by seeding primary uterine cells and MSCs on decellularized uterine matrix (Miyazaki and Maruyama, 2014), was less than the group implanted with SF-BC loaded with rhSDF-1α (7.00 ± 0.99) in this study (table3). Differences in the pregnancy outcomes amongst the three groups illustrated that SF-BC loaded with rhSDF-1α could initiate functional uterine regeneration that led to an improved pregnancy rate.

It has been reported that SDF-1α can promote the regeneration of various tissues by recruiting endogenous cells to the injury site, and facilitating the formation of blood vessels via recruitment of endothelial cells (Imitola *et al.*, 2004; Lau and Wang, 2011; Ho *et al.*, 2012; Gil *et al.*, 2013; Sun *et al.*, 2013; Song *et al.*, 2014). In this study, SDF-1 released from SF-BC may promote the migration of adjacent uterine cells to the injury site to take part in repair and regeneration of the uterus. Hence, the better pregnancy outcomes observed in this study may be due to the following mechanisms: 1) SDF-1 promotes vascularization. 2) and more mature endometrium formation.

SDF-1 promotes vascularization: In humans, uterine spiral arteries perfuse the inter-villous space, which enable the exchange of nutrients and oxygen between mother and fetus (Lima *et al.*, 2014). With decrease in vasodilation of uterine arteries in mice, fetal growth would be restricted, implying that uterine arteries are crucial to fetal growth by providing enough blood supply to the fetus (Gokina *et al.*, 2013). In our study, the number of arteries at the injury site of the uterus implanted with SF-BC loaded with rhSDF-1α was more than that in the uterus of the injury alone group at 30 and 90 days post-surgery. This may contribute to better pregnancy outcomes in the uterus implanted with SF-BC loaded with rhSDF-1α.

SDF-1 promotes more mature endometrium formation: Uterine glands are essential for fetus development in early pregnancy, which provide nutrients, growth factors, immune-regulatory proteins (Burton *et al.*, 2007) and some transport carriers (Hempstock *et al.*, 2004). In our study, there were more uterine glands in the regenerated endometrium of the SF-BC loaded with rhSDF-1α group. More uterine glands at the injury site of the uterus in the SF-BC loaded with rhSDF-1αgroup may lead to better pregnancy outcomes than the injury alone group, by providing more nutrients, like carbohydrates, glycoproteins and lipids to the embryo prior to its implantation into the endometrium (Burton *et al.*, 2007), more growth factors, such as LIF, VEGF, TNF-μ that promote the formation of utero-placental arteries and villous tree, more immune-regulatory proteins, like glycodelin A and MUC1 to facilitate embryo implantation (Burton *et al.*, 2007), and more transport carriers, such as TPP and lactoferrin to boost the immune tolerance of the feto-placenta and endometrium (Hempstock *et al.*, 2004).

## 5. Conclusion

In this study, we demonstrated that the SF-BC membrane possessed good physical, chemical and biocompatibility properties *in vitro*. The *in vivo* study showed that the incorporation of rhSDF-1α within the SF-BC membrane promoted regeneration of full-thickness uterine injury, and also improved the pregnancy outcome of the damaged uterus. The results thus suggest that SF-BC loaded with rhSDF-1α has good potential in future clinical applications for the repair of uterine injury.

## Acknowledgements

This work was supported by the National High Technology Research and Development Program of China (2016YFC1100401) (863 Program, 2015AA020302), the National Natural Science Foundation of China (CN) (81270682, 81300454), the Key Scientific and Technological Innovation Team of Zhejiang Province (2013TD11), China Postdoctoral Science Foundation (2015M571887), Zhejiang Medical and Health Science and Technology plan project (2013KYB080), and the Fundamental Research Funds for the Central Universities (2014QNA7016), and the Zhejiang Provincial Natural Science Foundation of China (LY14C100003).

## Author contributions Statement

H.X.C.: cell culture, animal experiments, acquisition of data, data analysis and interpretation, manuscript writing; B.B.W.: cell culture, animal experiments, acquisition of data, manuscript writing; Y.X.L.: animal experiments, manuscript writing; Y.L.: acquisition of data; L.B.S.: animal experiments; L.G.: acquisition of data; Y.X.X.: sampling of clinical samples; B.C.H.: manuscript writing; H.L.W.: sampling of clinical samples; H.W.O.Y.: conception and design; Z.H.Z.: preparation of biomaterials, manuscript writing; X.H.Z.: conception and design, manuscript writing;

## Disclosure of Potential Conflicts of Interest

The authors indicate no potential conflict of interest.

## Reference

Ajisawa, A. 1998 Dissolution of silk fibroin with calcium chloride-ethanol aqueous solution. J Seric Sci 67: 91–94.

Burton, G J, E Jauniaux,D S Charnock-Jones. 2007 Human Early Placental Development: Potential Roles of the Endometrial Glands. Placenta28, Supplement: S64–S69.

Ding, L, X a Li, H Sun, J Su, N Lin, B Péault, T Song, J Yang, J Dai,Y Hu. 2014 Transplantation of bone marrow mesenchymal stem cells on collagen scaffolds for the functional regeneration of injured rat uterus. Biomaterials35: 4888–4900.

Gil, M, M Seshadri, M P Komorowski, S I Abrams,D Kozbor. 2013 Targeting CXCL12/CXCR4 signaling with oncolytic virotherapy disrupts tumor vasculature and inhibits breast cancer metastases. Proceedings of the National Academy of Sciences 110: E1291–E1300.

Gokina, N I, S-L Chan, A C Chapman, K Oppenheimer, T L Jetton,M J Cipolla. 2013 Inhibition of PPARγ during rat pregnancy causes intrauterine growth restriction and attenuation of uterine vasodilation. Frontiers in Physiology4: 184.

Gu, Y, J Zhu, C Xue, Z Li, F Ding, Y Yang,X Gu. 2014 Chitosan/silk fibroin-based, Schwann cell-derived extracellular matrix-modified scaffolds for bridging rat sciatic nerve gaps. Biomaterials35: 2253–2263.

Hempstock, J, T Cindrova-Davies, E Jauniaux,G J Burton. 2004 Endometrial glands as a source of nutrients, growth factors and cytokines during the first trimester of human pregnancy: a morphological and immunohistochemical study. Reprod Biol Endocrinol2: 1–14.

Ho, T K, X Shiwen, D Abraham, J Tsui,D Baker. 2012 Stromal-cell-derived factor-1 (SDF-1)/CXCL12 as potential target of therapeutic angiogenesis in critical leg ischaemia. Cardiology research and practice2012.

Imitola, J, K Raddassi, K I Park, F-J Mueller, M Nieto, Y D Teng, D Frenkel, J Li, R L Sidman, C A Walsh, E Y Snyder,S J Khoury. 2004 Directed migration of neural stem cells to sites of CNS injury by the stromal cell-derived factor 1μ/CXC chemokine receptor 4 pathway. Proceedings of the National Academy of Sciences 101: 18117–18122.

Kim, J, S W Kim, S Park, K T Lim, H Seonwoo, Y Kim, B H Hong, Y-H Choung,J H Chung. 2013 Bacterial Cellulose Nanofibrillar Patch as a Wound Healing Platform of Tympanic Membrane Perforation. Advanced Healthcare Materials2: 1525–1531.

Lau, T T,D-A Wang. 2011 Stromal cell-derived factor-1 (SDF-1): homing factor for engineered regenerative medicine. Expert Opinion on Biological Therapy11: 189–197.

Lin N, X Li, T Song, J Wang, K Meng, J Yang, X Hou, J Dai, Y Hu. 2012 The effect of collagen-binding vascular endothelial growth factor on the remodeling of scarred rat uterus following full-thickness injury.Biomaterials 33:1801–1807.

Lindsay S. Wray, X Hu, J Gallego, I Georgakoudi, F. G. Omenetto, D Schmidt, D L. Kaplan. 2011 Effect of Processing on Silk-Based Biomaterials: Reproducibility and Biocompatibility. J Biomed Mater Res B Appl Biomater99: 89–101.

Lima, P D A, J Zhang, C Dunk, S J Lye,B Anne Croy. 2014 Leukocyte driven-decidual angiogenesis in early pregnancy. Cell Mol Immunol11: 522–537.

Micili, S, A Goker, O Sayin, P Akokay,B Ergur. 2013 The effect of lipoic acid on wound healing in a full thickness uterine injury model in rats. Journal of Molecular Histology44: 339–345.

Miyazaki, K,T Maruyama. 2014 Partial regeneration and reconstruction of the rat uterus through recellularization of a decellularized uterine matrix. Biomaterials35: 8791–8800.

Nakamura, Y, H Ishikawa, K Kawai, Y Tabata,S Suzuki. 2013 Enhanced wound healing by topical administration of mesenchymal stem cells transfected with stromal cell-derived factor-1. Biomaterials34: 9393–9400.

Shen, W, X Chen, J Chen, Z Yin, B C Heng, W Chen,H-W Ouyang. 2010 The effect of incorporation of exogenous stromal cell-derived factor-1 alpha within a knitted silk-collagen sponge scaffold on tendon regeneration. Biomaterials31: 7239–7249.

Son, B-R, L A Marquez-Curtis, M Kucia, M Wysoczynski, A R Turner, J Ratajczak, M Z Ratajczak,A Janowska-Wieczorek. 2006 Migration of Bone Marrow and Cord Blood Mesenchymal Stem Cells In Vitro Is Regulated by Stromal-Derived Factor-1-CXCR4 and Hepatocyte Growth Factor-c-met Axes and Involves Matrix Metalloproteinases. STEM CELLS24: 1254–1264.

Song, M, H Jang, J Lee, J H Kim, S H Kim, K Sun,Y Park. 2014 Regeneration of chronic myocardial infarction by injectable hydrogels containing stem cell homing factor SDF-1 and angiogenic peptide Ac-SDKP. Biomaterials35: 2436–2445.

Sun, X, C Charbonneau, L Wei, W Yang, Q Chen, R M Terek. 2013 CXCR4-Targeted Therapy Inhibits VEGF Expression and Chondrosarcoma Angiogenesis and Metastasis. Molecular Cancer Therapeutics12: 1163–1170.

Taveau, J W, M Tartaglia, D Buchannan, B Smith, G Koenig, K Thomfohrde, B Stouch, S Jeck,C H Greene. 2004 Regeneration of Uterine Horn Using Porcine Small Intestinal Submucosa Grafts in Rabbits. Journal of Investigative Surgery17: 81–92.

Tu, D D, Y G Chung, E S Gil, A Seth, D Franck, V Cristofaro, M P Sullivan, D Di Vizio, P Gomez Iii, R M Adam, D L Kaplan, C R Estrada Jr,J R Mauney. 2013 Bladder tissue regeneration using acellular bi-layer silk scaffolds in a large animal model of augmentation cystoplasty. Biomaterials34: 8681–8689.

Xu, F, J Li, V Jain, R S Tu, Q Huang,V Nanda. 2012 Compositional Control of Higher Order Assembly Using Synthetic Collagen Peptides. Journal of the American Chemical Society134: 47–50.

Yan, S, Q Zhang, J Wang, Y Liu, S Lu, M Li,D L Kaplan. 2013 Silk fibroin/chondroitin sulfate/hyaluronic acid ternary scaffolds for dermal tissue reconstruction. Acta Biomaterialia9: 6771–6782.

Zhang, W, J Chen, J Tao, Y Jiang, C Hu, L Huang, J Ji,H W Ouyang. 2013 The use of type 1 collagen scaffold containing stromal cell-derived factor-1 to create a matrix environment conducive to partial-thickness cartilage defects repair. Biomaterials34: 713–723.

Zhu, Z,H, Y H Dong, X Zhao,X H Zhou. 2015 Preparation of Regenerated Silk Fibroin/Bacterial Cellulose Com posite Membrane and Optimization of Fermentation Cultural Conditions. Science of Sericulture41: 0940–0945.

